# Revealing trajectories of multi-modal voxel-level changes in neurodegenerative diseases using latent event mapping

**DOI:** 10.64898/2026.06.07.730710

**Authors:** Sanduni Pinnawala, Annabelle Hartanto, Misha Jairamani, Ivor J.A. Simpson, Peter A. Wijeratne

## Abstract

Neurodegenerative diseases are driven by pathological mechanisms that can be indirectly measured in vivo using multi-modal neuroimaging. However, current computational methods that aim to reconstruct trajectories of voxel-level changes in the brain are either not computationally scalable or fully interpretable, limiting their ability to reveal associations between disease progression and underlying mechanisms. Here we introduce Latent Event Mapping (LEMING), a generative unsupervised modelling technique that learns a latent map of disease events along a common pseudo-timeline of events. We apply LEMING to amyloid PET and structural MRI data from the Alzheimer’s Disease Neuroimaging Initiative to reveal the first voxel-level trajectories of events in Alzheimer’s disease. Notably, we show how LEMING can provide new insights into progression-dependent disease mechanisms. We find that acetylcholine receptor density is significantly positively associated with both late-stage amyloid and atrophy events, suggesting that either these receptors are targeted later in disease progression, or that amyloid does not play an active role. This has strong implications for therapeutics that target acetylcholine receptors, particularly for early-stage intervention strategies.

## 1 Introduction

The pathological hallmarks of neurodegenerative diseases, such as Alzheimer’s disease (AD), accumulate silently for years before any clinical symptom emerges [1–3], yet nearly every therapeutic intervention occurs after this window has closed [2, 4, 5]. This presymptomatic period is presumably where disease-modifying intervention is most effective, and where our understanding of disease aetiology breaks down. Neurodegeneration is not a single process; molecular changes, such as amyloid accumulation, and structural changes, such as cortical atrophy, unfold in parallel, each with its own trajectory, each likely interacting with the other [1, 6–8]. But the order in which these changes occur together, across modalities and not in isolation, remains an open question. Poor understanding of this chain of events can lead to inconsistent biological narratives and trial designs that risk missing the optimal intervention time [4, 9, 10].

Disease progression models (DPMs) have emerged as a principled computational approach for reconstructing the temporal ordering of pathological events directly from observational data, without requiring extensive longitudinal follow-up. By pooling cross-sectional cohort data, these models infer a latent disease timeline, placing individuals along a common progression axis and estimating when each biomarker departs from normality [6, 11]. DPMs can be characterised into two broad classes: continuous DPMs, which estimate smooth biomarker trajectories along a latent disease axis [12– 17], and discrete DPMs, which infer the most probable ordering of biomarker events across a cohort [18–23]. Discrete models have continued to evolve because they yield directly interpretable disease cascades and remain robust in relatively small datasets (N ∼ 100-1000).

The event-based model (EBM) [18] established the core formulation: given cross-sectional biomarker measurements, infer the most probable sequence in which biomarkers become abnormal. Subsequent work extended the method to model disease subtypes with distinct progression sequences [20]. However, EBMs have operated largely on regional summary statistics, discarding the rich spatial structure present in the imaging data [11, 20]. More recently, the variational event-based model [24] reformulated event-based inference using optimal transport to achieve fast, scalable, inference of probabilistic event maps at the pixel level, treating each imaging feature as an independent event in a high-dimensional cascade. The closest comparative model is DIVE [25], which is a continuous DPM that operates on voxelwise imaging measures, clustering them by shared temporal dynamics and estimating continuous trajectories for each spatial cluster. DIVE requires longitudinal data and carries a substantially greater computational cost than the variational event-based model. Deep learning approaches such as variational autoencoders [26] have similarly been applied at high spatial resolution, but are essentially black boxes - their latent representations do not directly map onto meaningful event sequences, and demand orders of magnitude more data and compute.

Here we introduce latent event mapping (LEMING), a novel generative unsupervised modelling technique based on [24] for learning multi-modal voxel-level latent event mappings and their association with underlying disease mechanisms. Applied to amyloid PET and structural MRI-derived tensor-based morphometry data from the Alzheimer’s Disease Neuroimaging Initiative (ADNI), we demonstrate cross-modality tissue-level MRI-PET event timelines in AD. We extend the model to include cognitive biomarkers to reveal the relative positioning of voxel-level imaging changes alongside cognitive changes. We demonstrate fine-grained individual-level staging for early detection. Finally, we show associations between tissue-level atrophy, amyloid concentration, and cell-scale data, grounding tissue-level patterns in cell-scale biology and providing new insights into underlying disease mechanisms.

## 2 Results

### 2.1 Cross-modality tissue-level MRI-PET event timelines in AD

We peform LEMING on voxel-level amyloid PET (SUVr) and structural MRI-derived tensor-based morphometry (TBM) maps. Figure 1 shows the joint voxel-level event map, assigning each voxel a position along the common disease sequence separately for each modality. The event map reveals that amyloid accumulation, measured by SUVr, precedes structural atrophy, measured by TBM, across the brain. Early SUVr events are concentrated in (regions), consistent with known patterns of early amyloid deposition in AD [27]. Structural atrophy follows later on the timeline, with early TBM events localised to (regions), reflecting established patterns [28].

**Fig. 1:**
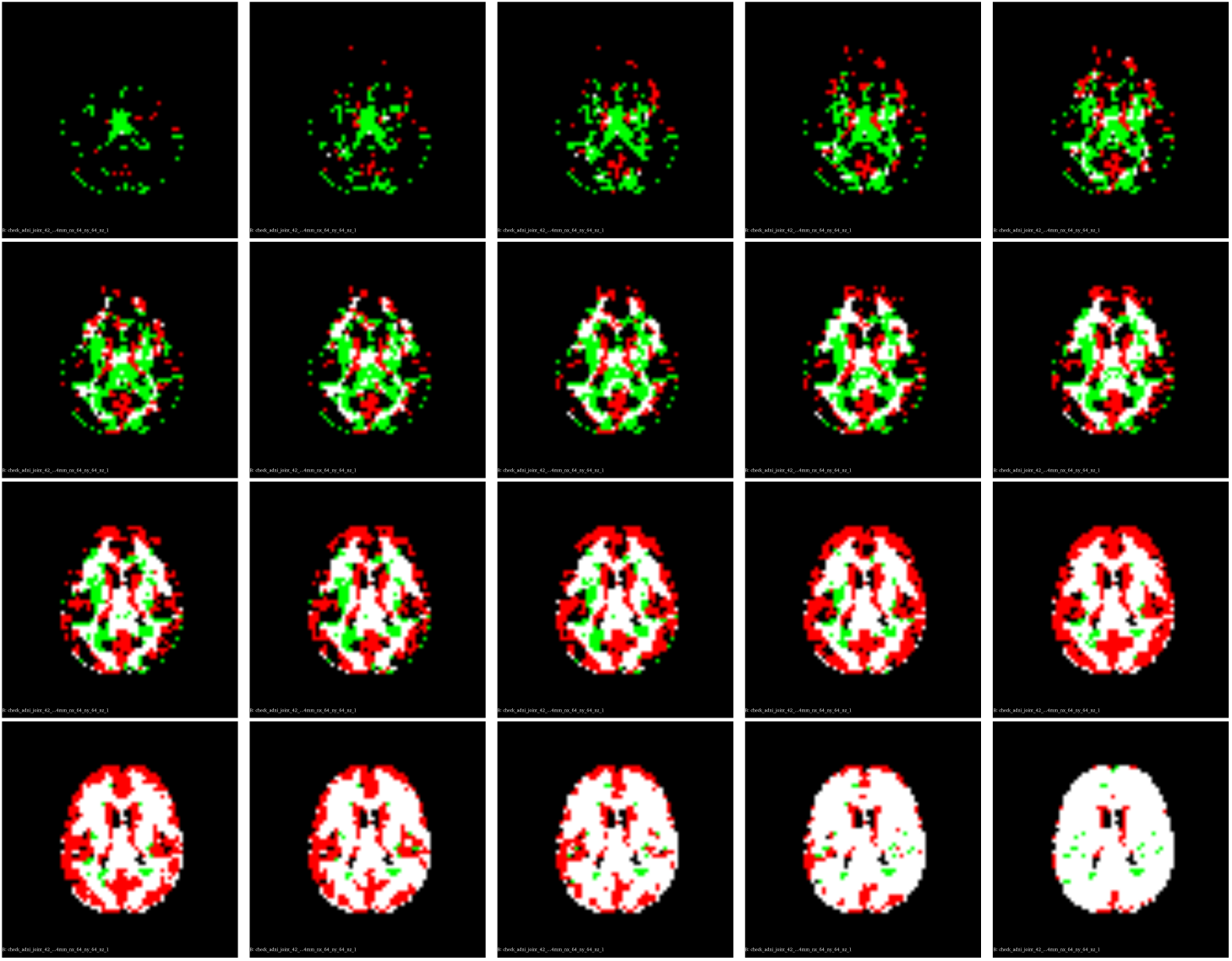
Multi-modal PET-MRI voxel-level disease progression sequence in AD obtained by LEMING. Green pixels are amyloid events; red pixels are atrophy events; white pixels are where both atrophy and amyloid events have occurred. The figure shows 20 events at uniform steps of 100 across the total of 2097, with the top left figure corresponding to event 100 (the first 100 events have occurred) and the bottom right to event 2000. Images were made using 3D Slicer (https://www.slicer.org/).

### 2.2 Associations between tissue-level atrophy, amyloid concentration and cell-scale mechanisms

We find the highest / lowest ACT density is at latest / earliest amyloid PET events (Spearman correlation = 0.25; p-value = 1E-16). We find a similar but weaker correlation with atrophy events (Spearman correlation = 0.1; p-value = 2E-3).

### 2.3 Mixed cognitive-imaging timelines

Imaging and cognitive changes follow a consistent sequence, with amyloid and atrophy events located centrally in the brain occur before downstream cognitive events. Notably, almost all atrophy events occur before MMSE becomes abnormal.

### 2.4 Fine-grained individual-level staging for early detection

LEMING supports individual-level disease staging by assigning individuals a stage corresponding to the most likely position along the event sequence, given their data. We find that the distributions of CN, MCI and AD groups is as expected (CN mean stage = 478; standard deviation = 661; AD mean stage = 1499; standard deviation = 556; MCI mean stage = 887; standard deviation = 786).

## 3 Discussion

Here we introduce LEMING, a novel generative unsupervised modelling technique for extracting voxel-level trajectories from multi-modal data. We used LEMING to reconstruct a discrete progression sequence across amyloid PET and structural MRI voxels from cross-sectional data.

The central finding of this study is that the pseudo-temporal ordering of disease events in AD is recoverable at the voxel level, across amyloid PET and structural MRI modalities, without the imposition of predefined regions of interest (Figure 1). The inferred sequence of events reveals a clear temporal pattern between amyloid accumulation and structural atrophy; amyloid preceding atrophy, propagating outward from sub-cortical regions before spreading across the cortex. This ordering is consistent with the amyloid-first hypothesis [27]. The data-driven recovery of this sequence is not merely confirmatory of an established biological narrative; it also validates LEMING as a credible basis for downstream analysis.

LEMING enables cross-comparisons between structural atrophy, amyloid deposition, and normative maps of neurotransmitter receptor densities (Figure 2). The spatial ordering of structural atrophy events is significantly associated with cortical M1 acetylcholine (ACT) receptor density, estimated from normative PET maps. Regions with higher ACT density tend to be affected later in the amyloid sequence, with receptor density increasing across the inferred timeline. This pattern is at odds with the hypothesis that cholinergic function modulates regional vulnerability to tissue loss. Similarly, ACT receptor density shows a weaker but still significant relationship with the spatial ordering of atrophy events. Taken together, these findings suggest that either ACT receptors are targeted later in disease progression, or that amyloid does not play an active role. This has strong implications for therapeutics that target acetylcholine receptors, particularly for early-stage intervention strategies.

**Fig. 2:**
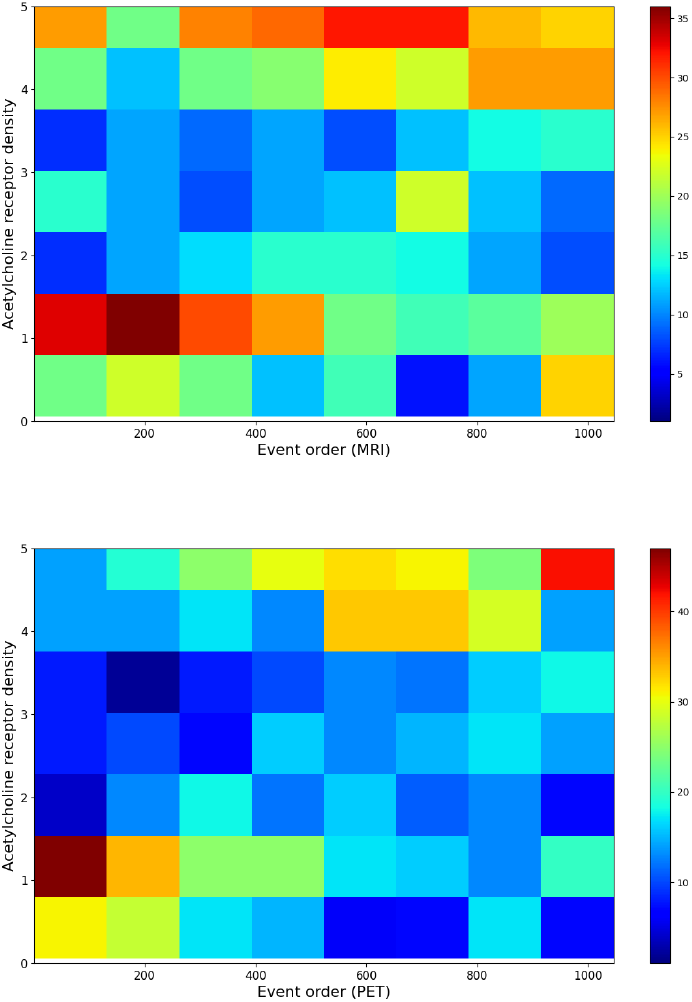
Heatmap (joint probability) of acetylcholine receptor density and LEMING event order. Top: atrophy events from MRI. Bottom: amyloid events from PET. The acetylcholine (ACT) receptor density map was obtained using the [neuromaps toolbox](https://github.com/netneurolab/neuromaps) [29].

The relative position of imaging and cognitive events reinforces this interpretation (Figure 3). LEMING identifies an extended presymptomatic window in which tissue-level pathological changes are already present while clinical cognitive measures remain largely unchanged. Existing EBMs provide only a coarse description of this phase, limiting their ability to obtain insights into tissue-level disease pathology and downstream cognitive changes.

**Fig. 3:**
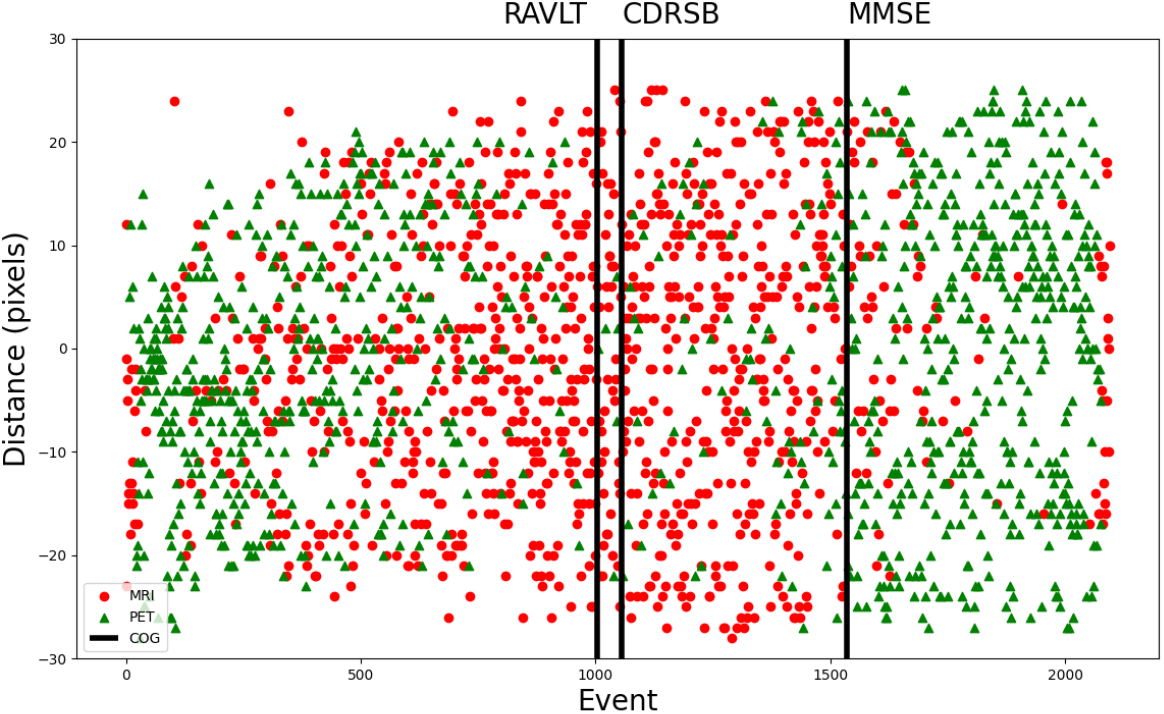
Multi-modal PET, MRI and cognitive events in AD. For imaging events, the vertical axis shows the distance from the centre of the brain to the event location. Cognitive events, denoted by vertical lines, do not have a spatial component. The horizontal axis shows the event ordering obtained by LEMING.

**Fig. 4:**
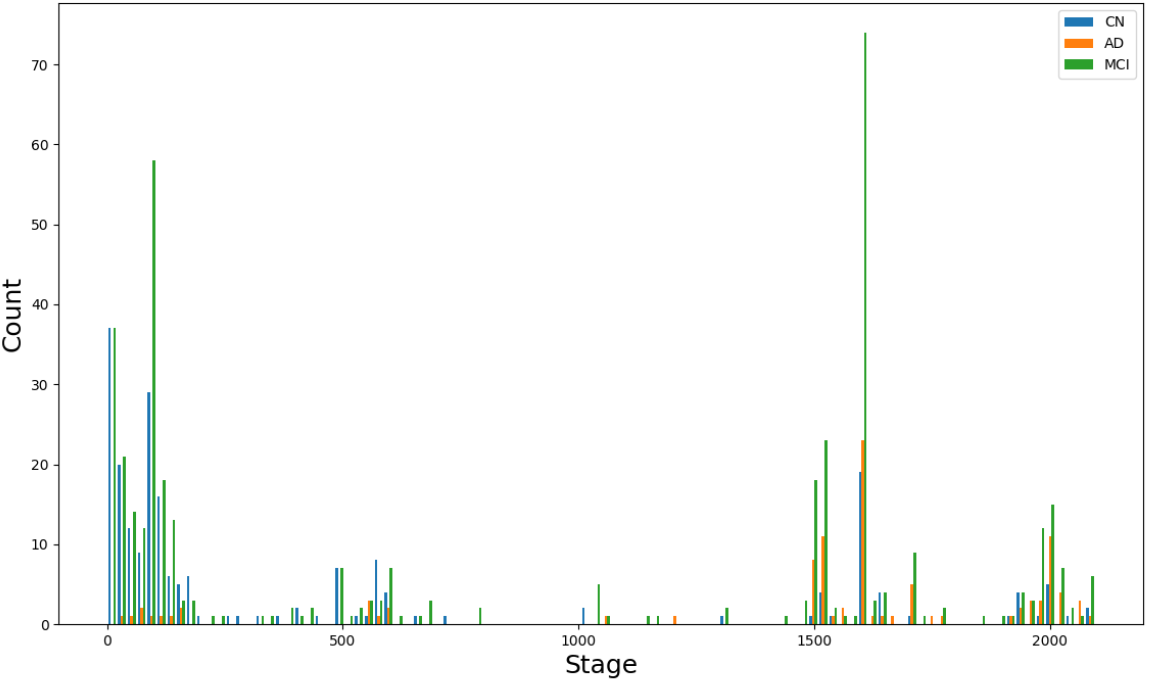
Individual-level staging from MRI-PET-COG LEMING. CN: cognitively normal; AD: Alzheimer’s disease; MCI: mild cognitive impairment.

As an image-based model operating directly at voxel resolution, LEMING is sensitive to preprocessing choices (e.g., inter-modal registration, smoothing), where errors in spatial alignment may propagate into event ordering and introduce systematic biases. Incorporating registration uncertainty directly into the inference framework would provide a principled means of mitigation. Although LEMING can accommodate longitudinal scans, we currently treat each visit as an independent snapshot, without explicitly modelling time intervals. This improves scalability but does not exploit the temporal information in longitudinal data; future extensions could integrate temporal event-based modelling [30] to link LEMING’s pseudo-temporal axis to absolute timescales, using the available follow-up intervals. The current formulation also assumes a single group-level progression sequence - a standard EBM assumption that may not fully capture the pathological heterogeneity of AD. Incorporating SuStaIn-like subtyping variants could recover distinct progression patterns corresponding to known subtypes. LEMING further inherits the EBM assumption of monotonic, irreversible biomarker progression, which may be violated locally by artefacts such as noise and partial volume effects; future work will focus on relaxing this constraint. We emphasise that LEMING recovers pseudo-temporal orderings and associations, not causal mechanisms; any mechanistic hypotheses require independent validation.

## 4 Conclusion

We introduce LEMING, a generative unsupervised model for mapping the joint spatiotemporal unfolding of pathological processes at voxel-level across imaging modalities. Applied here to AD, it recovers a coherent voxelwise sequence of amyloid accumulation and structural atrophy along a common disease timeline. While demonstrated on amyloid PET and structural MRI in AD, LEMING is modality-agnostic and disease-agnostic, applicable wherever multiple imaging modalities capture distinct aspects of a shared pathological process. We anticipate its application across the spectrum of neurodegenerative disease, where disentangling the sequence of co-occurring pathologies remains a central open problem.

## 5 Methods

### 5.1 Data

We use data from the ADNI database. ADNI data are available through an access request. Up-to-date information is available at https://adni.loni.usc.edu/. We include 794 ADNI participants with both T1-weighted MRI and AV45 PET data. The demographics of the participants, obtained from the ADNIMERGE table, are presented in Table 1.

**Table 1:**
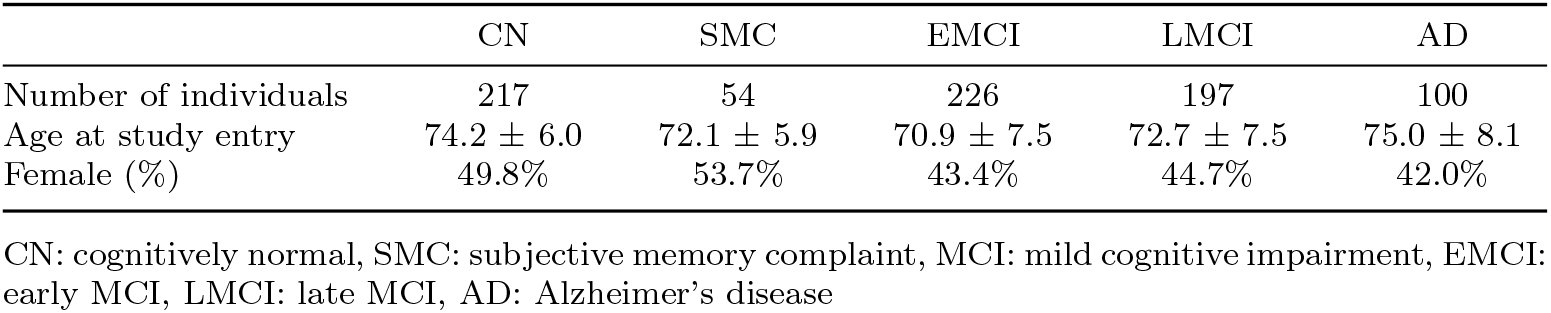
Demographics of the ADNI participants used in the study.

|  | CN | SMC | EMCI | LMCI | AD |
| --- | --- | --- | --- | --- | --- |
| Number of individuals | 217 | 54 | 226 | 197 | 100 |
| Age at study entry | $74.2 \pm 6.0$ | $72.1 \pm 5.9$ | $70.9 \pm 7.5$ | $72.7 \pm 7.5$ | $75.0 \pm 8.1$ |
| Female (%) | 49.8% | 53.7% | 43.4% | 44.7% | 42.0% |

CN: cognitively normal, SMC: subjective memory complaint, MCI: mild cognitive impairment, EMCI: early MCI, LMCI: late MCI, AD: Alzheimer’s disease

### 5.2 Code

Code for LEMING is available here: https://github.com/lililab-sussex/leming.

### 5.3 MRI processing

We download preprocessed T1-weighted MRI scans from ADNI. The ADNI MRI preprocessing pipeline includes scanner-level gradient nonlinearity correction using GradWarp and intensity nonuniformity correction using B1 and N3 correction.

We register the downloaded T1-weighted MRI scans to a common atlas space using the Advanced Normalisation Tools (ANTsPy v0.6.1) package. We use a two-stage registration process. First, each scan is registered to an intrasubject template using antsRegistrationSyN[sr], which includes rigid and deformable registration. Rigid registration corrects differences in head position and orientation between scans from the same participant. Deformable registration then accounts for local within-subject anatomical differences. Second, the intrasubject templates are registered to an intersubject atlas using ANTs SyN with affine and deformable registration. Affine registration accounts for global differences in position, rotation, scale and shear across participants. Deformable registration then improves local anatomical correspondence between each participant’s template and the atlas. This registration process places all participants in a common voxel coordinate system, allowing each voxel to approximately correspond to the same anatomical location across subjects.

We then derive cross-sectional tensor-based morphometry (TBM) maps from the nonlinear deformation fields. TBM quantifies voxel-wise structural morphology in atlas space from the deformation required to register an individual scan to the atlas. The Jacobian determinant of the deformation field measures local volume change, with values reflecting expansion or contraction relative to the atlas. We use the resulting TBM maps as MRI-derived voxel-level features for downstream event sequencing.

### 5.4 PET processing

We download preprocessed AV45 PET scans from ADNI. The ADNI PET preprocessing pipeline includes frame co-registration, frame averaging, standardisation of image and voxel size, and smoothing to a uniform 6mm resolution. Frame co-registration corrects within-scan motion, and the registered frames are averaged to generate a single static PET image for each visit. AV45 PET intensities are normalised to the cerebellar grey matter reference region to obtain standardised uptake value ratio (SUVr) maps. We use these SUVr maps as the PET measure of amyloid burden.

We then align each AV45 PET scan with the T1-weighted MRI scan from the nearest study time point. The PET scan and corresponding MRI scan are first transformed to the participant’s intrasubject template space. We then co-register the template-space PET image to the template-space MRI image, placing PET and MRI scans in a shared subject-specific anatomical space before atlas transformation. The co-registered PET image is then transformed to atlas space using the MRI-derived transformations.

This allows the SUVr maps and TBM maps to be analysed in a common voxel coordinate system. We modulate the PET SUVr maps using corresponding MRI-derived Jacobian determinant maps to account for volumetric changes introduced by transformation to atlas space. We use the resulting SUVr maps as PET-derived voxel-level features for downstream event sequencing.

We review all intermediate outputs at key stages of the processing pipeline to identify failures in registration and alignment. Scans failing quality control are excluded from modelling.

### 5.5 Mathematical model

We use the variational-event based model [24] as the base model to peform latent event mapping (LEMING). Here, an event represents the transition of a biomarker from a normal to an abnormal state. Each event corresponds to an abnormality in either a TBM feature or an SUVr feature at a given voxel. The model estimates the event sequence that best explains the observed pattern of biomarker abnormality across participants.

Formally, vEBM defines a generative latent variable model with observed data *Y* and latent variables *Z* = *{S, k, θ}*, where 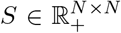 is a latent permutation matrix over *N* events, 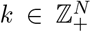 is the latent state of an individual, and *θ* denotes model parameters. The joint probability factorises as:

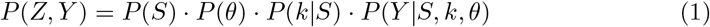

Each element of *S* defines an event – the transition of a feature from normal to abnormal relative to a reference distribution. For each feature *j*, normality and abnormality are characterised by univariate Gaussian distributions with mean 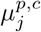, standard deviation 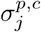, and mixture weight 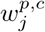, for patient (*p*) and control (*c*) distributions respectively. The discrete permutation *s ∈* Perm(*N*) is obtained from *S* over *N* events, and 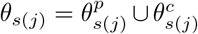 are the patient and control distribution parameters for feature *j* at position *s*(*j*) in the permutation. Under the assumptions of monotonic group-level progression and a consistent event ordering across the population, the model likelihood over *I* individuals and *J* features is [24]:

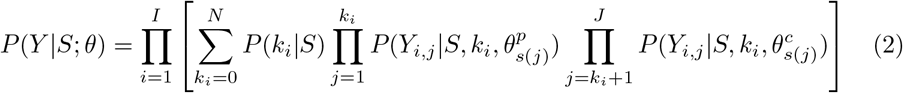

The model approximates the posterior over permutations using variational inference [24]. To enable differentiable permutation inference, the variational posterior is parameterised using the Gumbel-Sinkhorn distribution 𝒢 (*X, τ*), with *τ*, a temperature parameter controlling the trade-off between soft and hard permutations, and *ϵ*, a matrix of i.i.d. Gumbel noise:

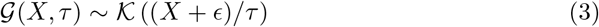

A uniform prior over permutations 𝒢 (*X* = 0, *τ*_prior_) is assumed, and the model is optimised by maximising the evidence lower bound (ELBO):

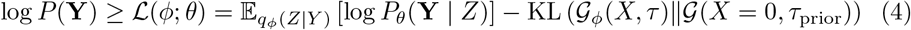

As such, LEMING represents disease progression as a continuous permutation of probabilities in a doubly stochastic matrix. This makes the mathematical object represented by LEMING fundamentally different to the event-based model: we now infer a matrix of continuous probabilities over events, rather than a discrete set of events; rather, the ordering emerges as a hard limit in LEMING. This unlocks gradient-based optimisation via variational inference, and also contains more information, such as event covariance.

### 5.6 Model hyperparameters

We train LEMING with the following settings: number of iterations = 200; step size = 1E-1; number of Sinkhorn iterations = 20; temperature = 1E3; temperature prior = 1; Gumbel scale = 0. For definitions of these hyperparameters see [24].

